# rehh 2.0: a reimplementation of the R package rehh to detect positive selection from haplotype structure

**DOI:** 10.1101/067629

**Authors:** Mathieu Gautier, Alexander Klassmann, Renaud Vitalis

## Abstract

Identifying genomic regions with unusually high local haplotype homozygosity represents a powerful strategy to characterize candidate genes responding to natural or artificial positive selection. To that end, statistics measuring the extent of haplotype homozygosity within (e.g., EHH, iHS) and between (Rsb or XP-EHH) populations have been proposed in the literature. The rehh package for R was previously developed to facilitate genome-wide scans of selection, based on the analysis of long-range haplotypes. However, its performance wasn’t sufficient to cope with the growing size of available data sets. Here we propose a major upgrade of the rehh package, which includes an improved processing of the input files, a faster algorithm to enumerate haplotypes, as well as multi-threading. As illustrated with the analysis of large human haplotype data sets, these improvements decrease the computation time by more than an order of magnitude. This new version of rehh will thus allow performing iHS-, Rsb- or XP-EHH-based scans on large data sets. The package rehh 2.0 is available from the CRAN repository (http://cran.r-project.org/web/packages/rehh/index.html) together with help files and a detailed manual.

## Introduction

Next-generation sequencing (NGS) technologies have deeply transformed the nature of polymorphism data. Although population geneticists were, until recently, limited by the amount of available data in a handful of presumably independent markers, they now have access to dense single nucleotide polymorphism (SNP) data in both model and non-model species (Davey *et al*, 2011). In those species where genome assemblies are available, the analysis of haplotype structure in a population has proved useful to detect recent positive selection (Sabeti *et al*, 2002). Consider neutral mutations appearing in a population: if, by chance, any of these increases in frequency after some time, then recombination should tend to break down linkage disequilibrium (LD) around it, thereby decreasing the length of haplotypes on which this mutation stands. Common variants are therefore expected to be old and standing on short haplotypes. If a mutation is selected for, however, it should expand in the population before recombination has time to break down the haplotype on which it occurred. A powerful strategy to characterize candidate genes responding to natural or artificial positive selection thus consists in identifying genomic regions with unusually high local haplotype homozygosity, relatively to neutral expectation (Sabeti *et al*, 2002).

For that purpose, Sabeti *et al* (2002) introduced a new metric, referred to as the extended haplotype homozygosity (EHH), which measures the decay of identity by descent, as function of distance, between randomly sampled chromosomes carrying a focal SNP. Tests of departure of EHH from neutral expectation were proposed, based on coalescent simulations of demographic history. Voight *et al* (2006) later introduced a test statistic (iHS) based on the standardized log-ratio of the integrals of the observed decay of EHH computed for the ancestral and the derived alleles at the focal SNP. Finally, cross-population statistics were proposed, to contrast EHH profiles between populations: XP-EHH (Sabeti *et al*, 2007) and Rsb (Tang *et al*, 2007). These haplotype-based methods of detecting selection have largely been applied on human data (Vitti *et al*, 2013), a wide range of livestock (see, e.g. Flori *et al*, 2014; Bosse *et al*, 2015; Barson *et al*, 2015) and plant species (see, e.g. Wang *et al*, 2014; Jin *et al*, 2016), and also non-model species (see, e.g. Roesti *et al*, 2015; Mueller *et al*, 2016).

A few years ago, we developed rehh (Gautier & Vitalis, 2012), a package for the statistical software package R (R Development Core Team, 2008), to detect recent positive selection from the analysis of long-range haplotypes. Since then, two alternative programs were released: `selscan` (Szpiech & Hernandez, 2014), which introduces multithreading to improve computational efficiency and `hapbin` (Maclean *et al*, 2015), which in addition to multithreading offers considerable gain in computation time thanks to a new computational approach based on a bitwise algorithm.

Here we propose a major upgrade of the rehh package (Gautier & Vitalis, 2012), which includes an improved algorithm to enumerate haplotypes, as well as multi-threading. These improvements decrease the computation time by more than an order of magnitude, as compared to the previous rehh version (1.13), which eases the analysis of big datasets.

Below we provide a brief overview of the statistics and tests available in rehh 2.0, and give a detailed worked example of the analysis of chromosome 2 in humans (HSA2), from HapMap samples CEU (Utah residents with Northern and Western European ancestry from the CEPH collection) and JPT+CHB (Japanese in Tokyo, Japan and Chinese from Beijing, China). We use this example as a guideline to use rehh2.0. We further show how rehh was improved since the previous version, and how it compares to the alternative programs `selscan` (Szpiech & Hernandez, 2014) and `hapbin` (Maclean *et al*, 2015).

## Overview of the EHH-based tests

### Within population tests

#### The allele-specific extended haplotype homozygosity: EHH (Sabeti *et al*, 2002)

At a focal SNP and for a given core allele (the ancestral or derived), the allele-specific extended haplotype homozygosity (EHH) is defined as the probability that two randomly chosen chromosomes (carrying the core allele considered) are identical by descent (IBD). IBD is assayed by computing homozygosity at all SNPs within an interval surrounding the core region (Sabeti *et al*, 2002). The EHH thus aims at measuring to which extent an extended haplotype is transmitted without recombination. In practice, the EHH (EHH_*as,t*_) of a tested core allele *a_s_* (*a_s_* = 1 or *a_s_* = 2) for a focal SNP s over the chromosome interval comprised between the core allele *a_s_* and the SNP *t* is computed as:

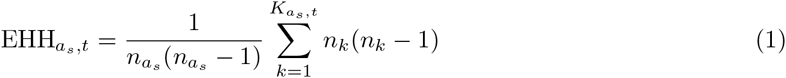

where *K_a_s_,t_* represents the number of different extended haplotypes (from SNP *s* to SNP *t*) carrying the core allele *a_s_, n_k_* is the number of the *k*th haplotype, and *n_a_s__* represents the number of haplotypes carrying the core allele *a_s_*, i.e., 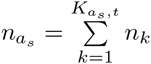.

#### The integrated (allele-specific) EHH: iHH (Voight *et al*, 2006)

By definition, irrespective of the allele considered, EHH starts at 1, and decays monotonically to 0 as one moves away from the focal SNP. For a given core allele, the integrated EHH (iHH) (Voight *et al*, 2006) is defined as the area under the EHH curve with respect to map position. In rehh (Gautier & Vitalis, 2012), this definite integral is computed using the trapezoidal rule. In practice, the integral is only computed for the regions of the curve above an arbitrarily small EHH value (e.g., EHH>0.05). In their seminal paper, Voight *et al* (2006) considered genetic distances and apply a penalty (proportional to physical distances) for successive SNPs separated by more than 20 kb. In addition, they did not compute iHH if any physical distance between a pair of neighboring SNPs was above 200 kb. We did not implement such an approach in rehh although this might easily be done by modifying the positions of the markers in SNP information input file. In addition, large gaps between successive SNPs (e.g., centromeres) might also be treated by splitting the chromosomes. For instance, when analyzing metacentric chromosomomes (e.g., HSA2), each chromosome arm may be considered separately in the analyses by assigning a different chromosome name (e.g., 2a and 2b) to the underlying SNPs.

#### The standardized ratio of core alleles iHH: iHS (Voight *et al*, 2006)

Let UniHS represent the log-ratio of the iHH for its ancestral (iHH_*a*_) and derived (iHH_*d*_) alleles:

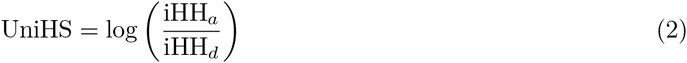

The iHS of a given focal SNP *s* (iHS(*s*)) is then defined following (Voight *et al*, 2006) as:

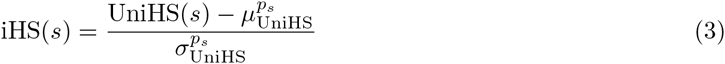

where 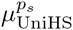 and 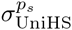 represent, respectively, the average and the standard deviation of the UniHS computed over all the SNPs with a derived allele frequency *p_s_* similar to that of the core SNP *s*. In practice, the derived allele frequencies are generally binned so that each bin is large enough (e.g., >10 SNPs) to obtain reliable estimates of 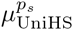 and 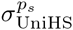. The iHS is constructed to have an approximately standard Gaussian distribution and to be comparable across SNPs regardless of their underlying allele frequencies. Hence, one may further transform iHS into p_iHS_ (Gautier & Naves, 2011):

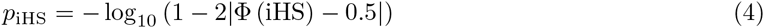

where Φ(*x*) represents the Gaussian cumulative distribution function. Assuming most of the genotyped SNPs behave neutrally (i.e., that the genome-wide empirical iHS distribution is a fair approximation of the neutral distribution), *p*_iHS_ may thus be interpreted as a two-sided *p*-value (in a – log_10_ scale) associated with the null hypothesis of selective neutrality.

### Pairwise-population tests

#### The site-specific extended haplotype homozygosity: EHHS (Tang *et al*, 2007; Sabeti *et al*, 2007)

At a focal SNP, the site-specific extended haplotype homozygosity (EHHS) is defined as the probability that two randomly chosen chromosomes are IBD at all SNPs within an interval surrounding the core region. EHHS might roughly be viewed as linear combination of the EHH’s for the two alternative alleles, with some weights depending on the corresponding allele frequencies. Two different EHHS estimators further referred to as EHHS^Sabeti^ and EHHS^Tang^ have been proposed by Sabeti *et al* (2007) and Tang *et al* (2007), respectively. For a focal SNP s over a chromosome interval extending to SNP *t*, these are computed as (using the same notation as above):

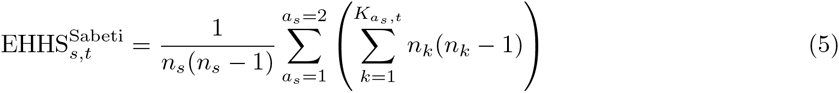

where 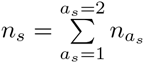 and

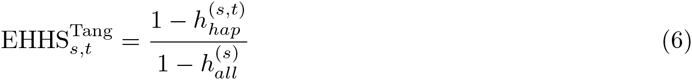

where:

- 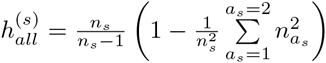 is an estimator of the focal SNP heterozygosity
- 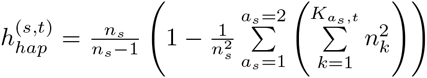 is an estimator of haplotype heterozygosity over the chro1mosome region interval extending from SNP *s* to SNP *t*.

#### The integrated EHHS: iES

As for the EHH (see above), EHHS starts at 1 and decays monotonically to 0 with increasing distance from the focal SNP. At a focal SNP, and in a similar fashion as the iHH, iES is defined as the integrated EHHS (Tang et al, 2007). Depending on the EHHS estimator considered, EHHS^Sabeti^ or EHHS^Tang^, two different iES estimators, that we further refer to as iES^Sabeti^ and iES^Tang^ can be computed.

#### The standardized ratios of pairwise population iES: XP-EHH (Sabeti *et al*, 2007) and Rsb (Tang *et al*, 2007)

For a given SNP *s*, let LRiES^Sabeti^(*s*) (respectively LRiES^Tang^(*s*)) represent the (unstandardized) log-ratio of the 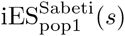 and 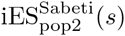 (respectively 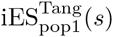 and 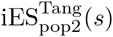) computed in two different populations:

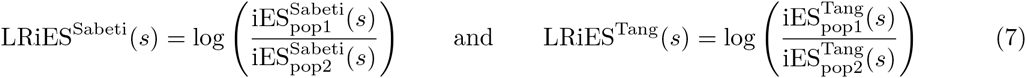

The XP-EHH (Sabeti *et al*, 2007) and the Rsb (Tang *et al*, 2007) for a given focal SNP are then standardized, as:

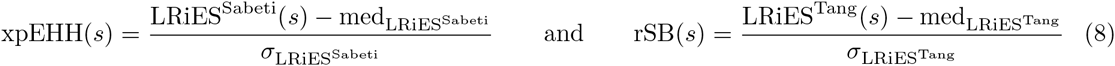

where med_LRiES^sab^_ (respectively med_LRiES^Tang^_) and *σ*_LRiES^sab^_ (respectively *σ*_LRiES^Tang^_) represent the median and standard deviation of the LRiES^Sabeti^(*s*) (respectively LRiES^Tang^(*s*)) computed over all the analyzed SNPs. As recommended by Tang *et al* (2007), the median is used instead of the mean because it is less sensitive to extreme data points. As for the iHS (see above), XP-EHH and Rsb are constructed to have an approximately standard Gaussian distribution. They may further be transformed into *p*_xpEHH_ or *p*_rSB_:

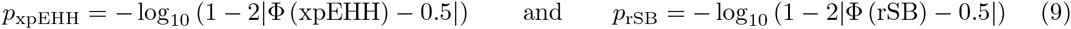

where Φ (*x*) represents the Gaussian cumulative distribution function. Assuming most of the genotyped SNPs behave neutrally (i.e., the genome-wide empirical distributions of XP-EHH and Rsb are fair approximations of their corresponding neutral distributions), *p*_xpEHH_ and *p*_rSB_ may thus be interpreted as a two-sided p-values (in a – log_10_ scale) associated with a null hypothesis of selective neutrality. Alternatively, one may also compute *p′*_xpEHH_ or *pȲ*_rSB_ as:

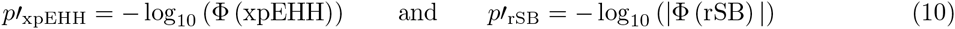

(see Gautier & Naves, 2011); *p′*_xpEHH_ and *p′*_rSB_ may then be interpreted as a one-sided *p*-values (in a – log10 scale) allowing the identification of those sites displaying outstandingly high EHHS in population *pop2* (represented in the denominator of the corresponding LRiES) relatively to the reference population (*pop*1).

## Material and Methods

### A new efficient algorithm to explore haplotype variability

In the previous version of rehh (1.13) the distribution of haplotype counts for the entire interval from the core SNP to the distance *x* was computed for each *x* independently, entailing repeatedly the same calculations. In the new version of rehh (2.0), the distribution of haplotype counts for the interval from the core SNP to the distance *x* is updated consecutively from the distribution of haplotype counts corresponding to the interval between the core SNP and *x* – 1. The new algorithm doesn’t affect the output, in particular, as in the previous version, all haplotypes carrying missing data are discarded from the computation of long-range homozygosity.

### Human haplotype data

Two HSA2 haplotype data sets were downloaded from the HAPMAP project (phase III) (The International HapMap3 Consortium, 2010) website (ftp://ftp.ncbi.nlm.nih.gov/hapmap/). They consisted of 236 haplotypes of 116,430 SNPs from the CEU and 342 haplotypes from the JPT+CHB populations, respectively. Further details about these data (including the phasing procedure) can be found on the HAPMAP website. For each SNP, the ancestral (and derived) allele was determined according to the Chimpanzee genome reference (using the *dbsnp_chimp_B36.gff* annotation file available at ftp://ftp.ncbi.nlm.nih.gov/hapmap/gbrowse/2010-08_phaseII+III/gff/). Such ancestral information is indeed required to carry out iHS-based tests (see above). As a result, 6,230 SNPs (5.35%) for which ancestral/derived states could not be unambiguously determined were discarded from further analyses leading to a total of 110,200 SNPs per analyzed haplotype.

### Computation

For comparison purposes, the different haplotype data sets were analyzed using the software packages rehh (both the previous version 1.13 and the new version 2.0), `selscan` (version 1.1.0b) (Szpiech & Hernandez, 2014) and `hapbin` (version 1.0.0) (Maclean *et al*, 2015). Default options were generally used except for the minimal threshold on the minor allele frequency (MAF) that was set to 0.01 for all programs. In addition, for the `selscan` program, both the window size around the core SNPs (`--ehh-win` option) and the maximum allowed gap in bp between two consecutive SNPs (`--max-gap` option) were set to 10^9^ (this was made to disallow these options that are not considered in other programs). Similarly, the `--max-extent` option was inactivated by setting `--max-extent=-1`. For the `hapbin` programs (i.e., `ihsbin` and `xpehhbin`), the EHH and EHHS cut-off values (defined to stop the calculation of unstandardized iHS and iES) were set to 0.05 (i.e., the default value in `selscan` and rehh). For all programs, the standardization of iHS was performed with allele frequency bins of 0.01, as controlled by the `freqbins` argument in the `ihh2ihs()` function of the rehh package, and the `bins` argument for the program `norm` of the `selscan` package and the program `ihsbin` of the `hapbin` package. The command lines used for the different programs, together with the corresponding input data files are provided in the Supplementary Materials.

Finally, for each analysis and parameter set, the estimation of computation times was averaged over ten independent runs. All analyses were run on a standard computer running under Linux Debian 8.5 and equipped with an Intel^®^ Xeon^®^ 6-core processor W3690 (3.46 GHz, 12M cache). Note that the Unix command `taskset` was used to control the number of working threads for the analyses with the `hapbin` programs (since neither the `ihsbin` nor the `xpehhbin` programs allow to chose the number of threads to be used).

## Results and Discussion

### Analysis of the human chromosome 2 data sets

For illustration purpose, we used rehh 2.0 to analyze two human data sets consisting of 236 and 342 haplotypes of 110,200 SNPs mapping to HSA2 that were sampled in the CEU and JPT+CHB populations respectively. The chromosome-wide scans of iHS for the CEU and the JPT+CHB populations, respectively, are plotted in Figure 1A. The most significant SNP map at position 136,503,121 bp for the CEU population (iHS=-5.35) and at position 111,506,728 bp for the JPT+CHB population (iHS=-4.92). The chromosome-wide scans of XP-EHH and Rsb, which contrast EHH profiles between the CEU and the JPT+CHB populations, are plotted in Figure 1B. The most significant SNP mapped at position 136,533,558 bp for Rsb-based test (Rsb=6.13) and at position 136,523,244 bp for the XP-EHH-based test (XP-EHH=5.59). For this latter SNP (mapping to region #7 as defined below), the haplotype bifurcation diagrams for the ancestral and derived alleles within the CEU population are plotted in (Figure 1C) and (Figure 1D), respectively, using the `bifurcation.diagram` function from the rehh package. Note the extent of haplotype homozygosity associated with the derived allele (Figure 1C), relatively to that associated with the ancestral allele (Figure 1D), which is consistent with the negative iHS measure at this SNP (iHS=-3.24).

**Figure 1.**
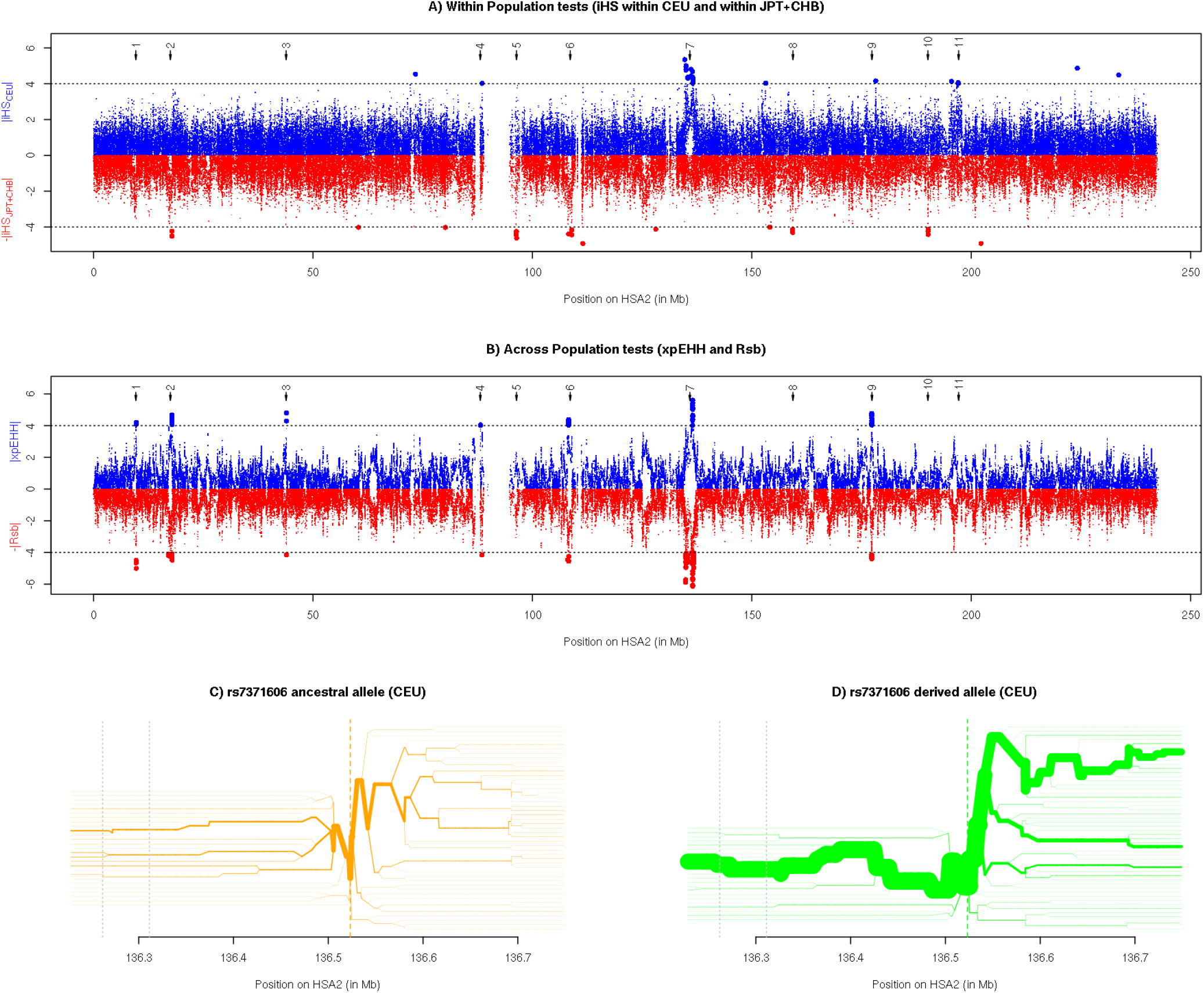
Analysis of the human chromosome 2 haplotype data sets (hg18 human genome assembly) for the CEU and JPT+CHB populations with rehh 2.0. A) Plot of iHS against physical distance, in the CEU (|iHS| in blue) and the JPT+CHB (— |iHS| in red) populations. B) Plot of XP-EHH (|XP-EHH| in blue) and Rsb (– |Rsb| in red) between the CEU and JPT+CHB populations. In A) and B), the horizontal dotted lines indicate the |iHS| significance threshold of 4 that was used to identify significant regions (see Table 1) and the arrows at the top of the graph indicate the mid-position of the significant regions described in Table 1). C) and D) Haplotype bifurcation diagrams drawn for the ancestral and derived allele, respectively, of the rs7377606 SNP in the CEU population (XP-EHH peak position of region #7 described in Table 1 and containing the LCT gene). In C) and D), the two grey vertical dotted lines delimit the LCT gene.

To further identify regions displaying strong footprints of selection, we split the HSA2 chromosome into 950 consecutive 500 kb-windows (with a 250 kb overlap). Windows with at least 2 SNPs displaying a statistic > 4 (in absolute value that roughly corresponds to a two-sided *p*-value< 10^-4^, see above) for at least one of the four test statistics were deemed significant. Significant overlapping windows were then merged, leading to a total of 11 regions harboring strong signals of selection, which characteristics are detailed in Table 1 (see also Figure 1). As expected, most of the regions identified in previously published genome-scans for samples with the same origin (Sabeti *et al*, 2007; Tang *et al*, 2007; Voight *et al*, 2006) overlap with the regions identified here (Table 1). For instance, regions #6 and #7 that are in the vicinity of the EDAR gene (under selection in Asian populations) and the LCT gene (under selection in European populations), respectively, have been extensively characterized in the literature (e.g., Peter *et al*, 2012). Interestingly, we detected more regions than previously reported in the aforementioned studies, most probably because our analyses are based on a larger dataset and different cut-off values. A more detailed description of the newly identified regions is however beyond the scope of the present article.

**Table 1.**
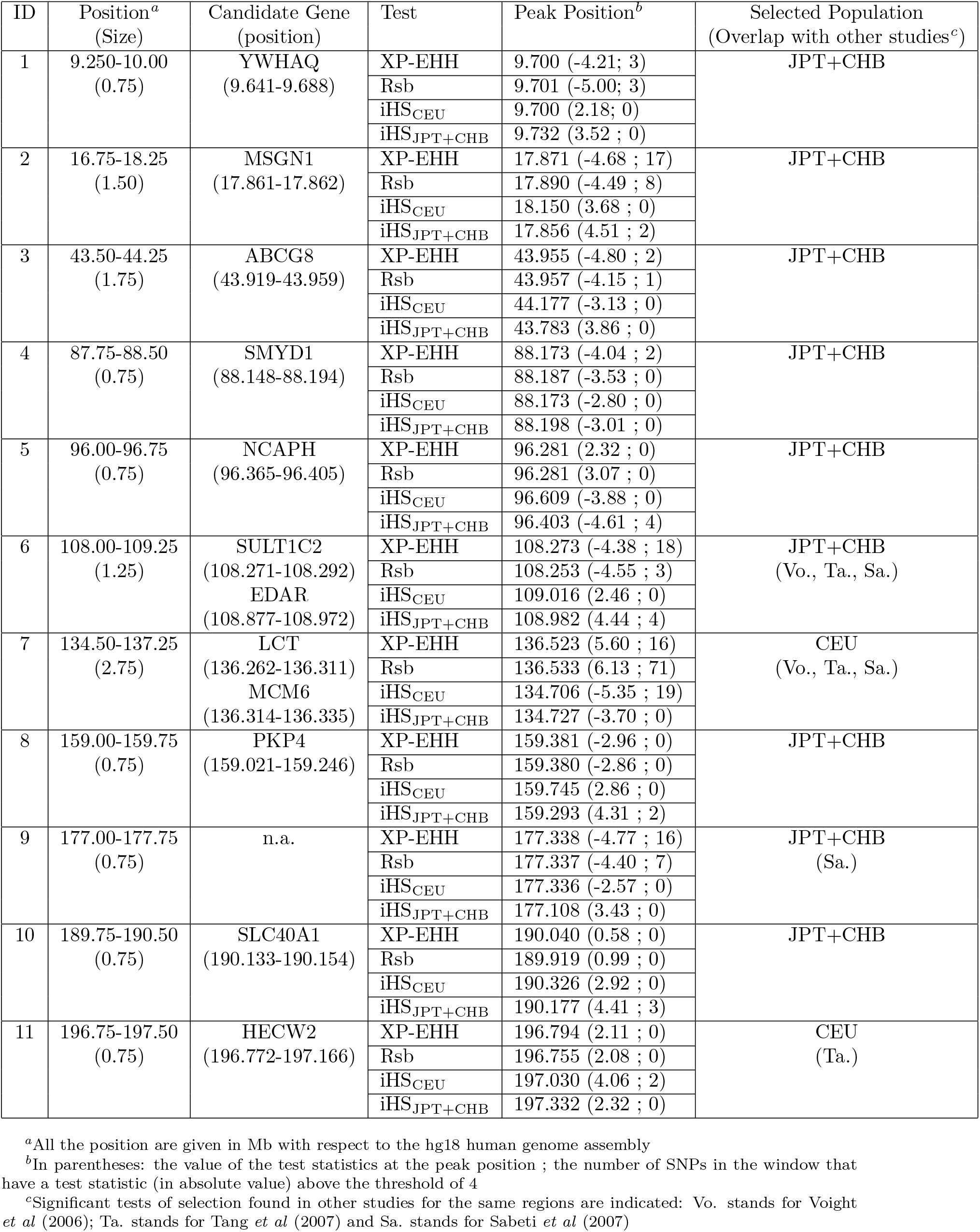
Regions of HSA2 harboring strong signals of selection.

| ID | Position<sup>a</sup><br>(Size) | Candidate Gene<br>(position) | Test | Peak Position<sup>b</sup> | Selected Population<br>(Overlap with other studies<sup>c</sup>) |
| --- | --- | --- | --- | --- | --- |
| 1 | 9.250-10.00<br>(0.75) | YWHAQ<br>(9.641-9.688) | XP-EHH | 9.700 (-4.21; 3) | JPT+CHB |
|  |  |  | Rsb | 9.701 (-5.00; 3) |  |
|  |  |  | iHS<sub>CEU</sub> | 9.700 (2.18; 0) |  |
|  |  |  | iHS<sub>JPT+CHB</sub> | 9.732 (3.52 ; 0) |  |
| 2 | 16.75-18.25<br>(1.50) | MSGN1<br>(17.861-17.862) | XP-EHH | 17.871 (-4.68 ; 17) | JPT+CHB |
|  |  |  | Rsb | 17.890 (-4.49 ; 8) |  |
|  |  |  | iHS<sub>CEU</sub> | 18.150 (3.68 ; 0) |  |
|  |  |  | iHS<sub>JPT+CHB</sub> | 17.856 (4.51 ; 2) |  |
| 3 | 43.50-44.25<br>(1.75) | ABCG8<br>(43.919-43.959) | XP-EHH | 43.955 (-4.80 ; 2) | JPT+CHB |
|  |  |  | Rsb | 43.957 (-4.15 ; 1) |  |
|  |  |  | iHS<sub>CEU</sub> | 44.177 (-3.13 ; 0) |  |
|  |  |  | iHS<sub>JPT+CHB</sub> | 43.783 (3.86 ; 0) |  |
| 4 | 87.75-88.50<br>(0.75) | SMYD1<br>(88.148-88.194) | XP-EHH | 88.173 (-4.04 ; 2) | JPT+CHB |
|  |  |  | Rsb | 88.187 (-3.53 ; 0) |  |
|  |  |  | iHS<sub>CEU</sub> | 88.173 (-2.80 ; 0) |  |
|  |  |  | iHS<sub>JPT+CHB</sub> | 88.198 (-3.01 ; 0) |  |
| 5 | 96.00-96.75<br>(0.75) | NCAPH<br>(96.365-96.405) | XP-EHH | 96.281 (2.32 ; 0) | JPT+CHB |
|  |  |  | Rsb | 96.281 (3.07 ; 0) |  |
|  |  |  | iHS<sub>CEU</sub> | 96.609 (-3.88 ; 0) |  |
|  |  |  | iHS<sub>JPT+CHB</sub> | 96.403 (-4.61 ; 4) |  |
| 6 | 108.00-109.25<br>(1.25) | SULT1C2<br>(108.271-108.292)<br>EDAR<br>(108.877-108.972) | XP-EHH | 108.273 (-4.38 ; 18) | JPT+CHB<br>(Vo., Ta., Sa.) |
|  |  |  | Rsb | 108.253 (-4.55 ; 3) |  |
|  |  |  | iHS<sub>CEU</sub> | 109.016 (2.46 ; 0) |  |
|  |  |  | iHS<sub>JPT+CHB</sub> | 108.982 (4.44 ; 4) |  |
| 7 | 134.50-137.25<br>(2.75) | LCT<br>(136.262-136.311)<br>MCM6<br>(136.314-136.335) | XP-EHH | 136.523 (5.60 ; 16) | CEU<br>(Vo., Ta., Sa.) |
|  |  |  | Rsb | 136.533 (6.13 ; 71) |  |
|  |  |  | iHS<sub>CEU</sub> | 134.706 (-5.35 ; 19) |  |
|  |  |  | iHS<sub>JPT+CHB</sub> | 134.727 (-3.70 ; 0) |  |
| 8 | 159.00-159.75<br>(0.75) | PKP4<br>(159.021-159.246) | XP-EHH | 159.381 (-2.96 ; 0) | JPT+CHB |
|  |  |  | Rsb | 159.380 (-2.86 ; 0) |  |
|  |  |  | iHS<sub>CEU</sub> | 159.745 (2.86 ; 0) |  |
|  |  |  | iHS<sub>JPT+CHB</sub> | 159.293 (4.31 ; 2) |  |
| 9 | 177.00-177.75<br>(0.75) | n.a. | XP-EHH | 177.338 (-4.77 ; 16) | JPT+CHB<br>(Sa.) |
|  |  |  | Rsb | 177.337 (-4.40 ; 7) |  |
|  |  |  | iHS<sub>CEU</sub> | 177.336 (-2.57 ; 0) |  |
|  |  |  | iHS<sub>JPT+CHB</sub> | 177.108 (3.43 ; 0) |  |
| 10 | 189.75-190.50<br>(0.75) | SLC40A1<br>(190.133-190.154) | XP-EHH | 190.040 (0.58 ; 0) | JPT+CHB |
|  |  |  | Rsb | 189.919 (0.99 ; 0) |  |
|  |  |  | iHS<sub>CEU</sub> | 190.326 (2.92 ; 0) |  |
|  |  |  | iHS<sub>JPT+CHB</sub> | 190.177 (4.41 ; 3) |  |
| 11 | 196.75-197.50<br>(0.75) | HECW2<br>(196.772-197.166) | XP-EHH | 196.794 (2.11 ; 0) | CEU<br>(Ta.) |
|  |  |  | Rsb | 196.755 (2.08 ; 0) |  |
|  |  |  | iHS<sub>CEU</sub> | 197.030 (4.06 ; 2) |  |
|  |  |  | iHS<sub>JPT+CHB</sub> | 197.332 (2.32 ; 0) |  |

<sup>a</sup>All the position are given in Mb with respect to the hg18 human genome assembly
<sup>b</sup>In parentheses: the value of the test statistics at the peak position ; the number of SNPs in the window that have a test statistic (in absolute value) above the threshold of 4
<sup>c</sup>Significant tests of selection found in other studies for the same regions are indicated: Vo. stands for Voight *et al* (2006); Ta. stands for Tang *et al* (2007) and Sa. stands for Sabeti *et al* (2007)

Note finally that XP-EHH- and Rsb-based scans gave consistent results, with the exception of the region in the vicinity of the LCT gene (#7 in Table 1 and Figure 1) where a double peak was observed with Rsb (consistent with the iHS profile within CEU) and a single peak with XP-EHH. The overall correlation between these statistics was equal to 0.843, which illustrates the close similarity of these two metrics.

### Comparing the performances of rehh 2.0 relatively to rehh 1.13, selscan and hapbin packages

The two CEU and JPT+CHB human data sets were further analyzed with rehh 1.13 to evaluate the gain in computation time resulting from the modifications introduced in version 2.0. Note that extensive tests were done during the development of version 2.0, to ensure that the same estimates were obtained with both versions. Only very marginal differences were however sometimes observed in the estimates of iES^Tang^. For instance, the correlation between the resulting Rsb computed across the CEU and JPT+CHB populations with version rehh 1.13 and rehh 2.0 was found equal to 0.999992 (instead of 1.0). This is actually due to the introduction of the computation of iES^Sabeti^ in version 2.0 to estimate XP-EHH. Indeed, we chose to define the same cut-off value for both statistics during the computation of the component variable EHHS (controlled with the option limehhs, set to 0.05 by default).

#### An improved processing of the input file

The first major modification introduced in rehh version 2.0 deals with the processing of input files (haplotype and SNP information files) using the function `data2haplohh`. Indeed our own experience with earlier versions of the package together with feedback from several users prompted us to optimize data import and to improve allele recoding, which was inefficient in version 1.x. Considering standard input haplotype file format (which is common to both versions), and with alleles encoded in the appropriate format ({0,1,2} for missing data, ancestral and derived alleles respectively), the new `data2haplohh` function is about 2.5 times faster than the previous one (see Table 2). In addition, the allele recoding option results in slightly better processing performances, and is no more prone to errors as in version 1.x. Finally, the new haplotype format (with haplotypes in columns), corresponding to the output file of the SHAPEIT phasing program (O’Connell *et al*, 2014), was found to be the most efficient to process (see Table 2).

**Table 2.**
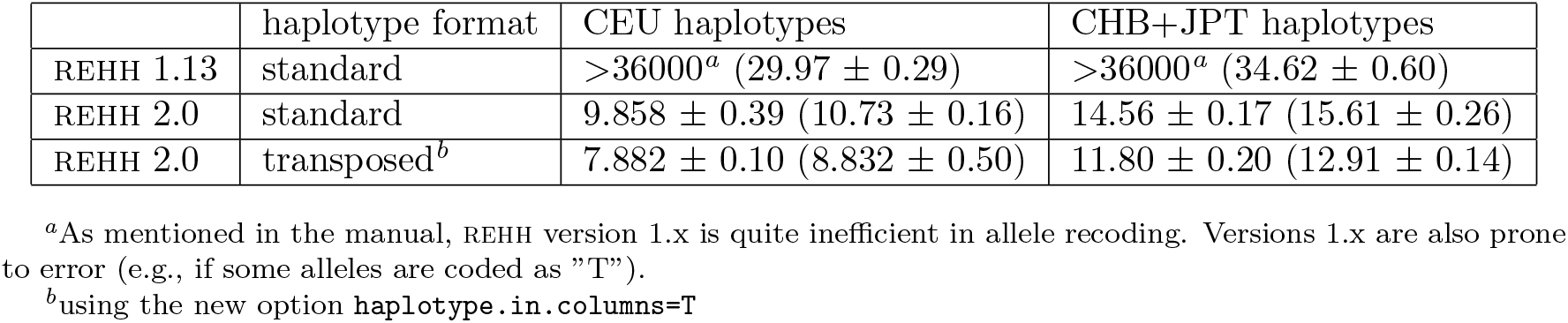
Comparison of the computation times (in seconds) required to process input data files with the *data2haplohh* function for the versions 1.13 and 2.0 of the rehh package. Two data sets consisting, respectively, of 236 and 342 haplotypes of 110,200 SNPs for the CEU and JPT+CHB populations were considered (see the main text). For each of these datasets, the table gives the average computation times (± standard deviation) across ten independent runs, either with or without (in parentheses) allele recoding (using the option `allele.recode`).

|  | haplotype format | CEU haplotypes | CHB+JPT haplotypes |
| --- | --- | --- | --- |
| REHH 1.13 | standard | >36000<sup>a</sup> (29.97 ± 0.29) | >36000<sup>a</sup> (34.62 ± 0.60) |
| REHH 2.0 | standard | 9.858 ± 0.39 (10.73 ± 0.16) | 14.56 ± 0.17 (15.61 ± 0.26) |
| REHH 2.0 | transposed<sup>b</sup> | 7.882 ± 0.10 (8.832 ± 0.50) | 11.80 ± 0.20 (12.91 ± 0.14) |

<sup>a</sup>As mentioned in the manual, REHH version 1.x is quite inefficient in allele recoding. Versions 1.x are also prone to error (e.g., if some alleles are coded as "T").
<sup>b</sup>using the new option `haplotype.in.columns=T`

With datasets of increasing complexity and size, such improvement in the processing of input files is critical to rehh users. Processing a data set as large as the JPT+CHB one (consisting of 342 haplotype with 110,200 SNPs) now takes less than 12 seconds. Note however that for this file a maximum of about 1 Gb RAM was used, for a net memory size change of 240 Mb. For larger data sets, RAM requirements may therefore be limiting for some computers.

#### A faster and parallel algorithm to explore haplotype variability

The second major modification introduced in rehh version 2.0 concerns the core algorithm that computes the distribution of haplotype counts, which underlies the calculation of all the metrics of interest (iHS, Rsb and XP-EHH). As shown in Table 3, this new algorithm allows to decrease the computation times by more than one order of magnitude, as compared to the algorithm implemented in rehh version 1.13. Hence, for the computation of iHS in the CEU population (respectively, the JPT+CHB population) on a single thread, computation times were 13.7 (respectively 21.8) times smaller on average. Interestingly, the computation time for the JPT+CHB dataset (which is approximately 1.34 times larger than the CEU one in terms of number of SNPs × number of haplotype) was only 1.09 times slower than for the latter. Conversely, the computation time was 1.73 times slower for JPT+CHB relatively to CEU with rehh version 1.13. Although a more detailed profiling of the algorithm would be required, these results suggest that computational burden is approximately linearly related to the data set complexity.

To further improve computational speed, haplotype structure is now performed using OpenMP parallelization across SNPs in genome-wide scans. Using four threads then lead to an additional decrease of about 3.5 times in computation times (see Table 3).

**Table 3.** Comparison of the time (in seconds) required to compute the different. EHH-based statistics for the versions 1.13 and 2.0 of the rehh, the `selscan` and the `hapbin` packages. For each analysis, the table gives the average computation time (± standard deviation) across ten independent runs. For each program, analyses were run either on a single thread or on four threads (except for rehh 1.13 version, which is not parallelised)

| program | #threads | iHS<sub>ceu</sub> | iHS<sub>chb+jpt</sub> | XP-EHH | Rsb | Total<sup>a</sup> |
| --- | --- | --- | --- | --- | --- | --- |
| REHH 1.13 | 1 | 1759 ± 29 | 3045 ± 31 | n.a. | 4803 ± 58 | 4805 ± 58 |
| REHH 2.0 | 1 | 128 ± 1.0 | 140 ± 2.1 | 268 ± 1.8 | 268 ± 1.8 | 269 ± 1.8 |
| | 4 | 37.8 ± 0.3 | 40.2 ± 0.3 | 77.1 ± 0.5 | 77.1 ± 0.5 | 78.5 ± 0.5 |
| **selscan** | 1 | 1237 ± 17 | 1503 ± 29 | 3833 ± 100 | n.a. | 6573 ± 86 |
| | 4 | 324 ± 6.5 | 391 ± 6.5 | 969 ± 5.6 | n.a. | 1684 ± 9.3 |
| **hapbin** | 1 | 17.6 ± 0.2 | 20.0 ± 0.1 | 47.4 ± 0.2 | n.a. | 85.0 ± 0.3 |
| | 4 | 5.68 ± 0.7 | 7.42 ± 0.1 | 13.2 ± 0.0 | n.a. | 26.2 ± 0.7 |

<sup>a</sup>In REHH the function **scanhh** computes iHH and iES simultaneously. It therefore needs to be run only once per haplotype data set. As a result, computing XP-EHH (and/or Rsb) requires almost no extra time, once iHS for the two populations has been computed.

Overall, the whole analysis of the HSA2 haplotype files used in this study took about 1.5 minutes (including the processing of input files) with rehh 2.0, and more than 1.3 hours with rehh 1.3. This corresponds to the computation of iHS within the CEU and within the JPT+CHB populations, as well as the computation of Rsb and XP-EHH.

### Comparing rehh 2.0 to the selscan and hapbin programs

Finally, we compared rehh 2.0 with `selscan` (Szpiech & Hernandez, 2014) and `hapbin` (Maclean *et al*, 2015), which were recently published. Both programs are written in C++ language and include parallelisation. Computation times for the different analyses, either on a single or four threads, are provided in Table 3. The new version of rehh outperforms `selscan` by about one order of magnitude. Moreover, running rehh on a single thread is still more than twice as fast as running `selscan` on four threads. It should also be noticed that running a full analysis consisting of the estimation of iHS within and XP-EHH between the CEU and JPT+CHB populations result in a significant additional burden with `selscan` (Table 3). Conversely, `hapbin` was found to be more than five times faster than rehh 2.0, most likely as a result of its more efficient algorithm to explore haplotype variability. Yet, given the small computation times achieved by both programs, rehh 2.0 remains competitive relative to `hapbin` for most practical applications.

Correlation between the estimated iHS and XP-EHH obtained with the different programs are given in Table 4. Estimates of XP-EHH were in almost perfect agreement among the different software packages. Similarly, estimates for iHS were almost the same between rehh 2.0 and `selscan` but slightly depart from those obtained with `hapbin`. Although we did not further investigate the origin of these discrepancies, this might probably be related to a different definition of haplotype homozygosity in `hapbin`, as compared to Sabeti *et al* (2007).

**Table 4.** Correlation between the estimated. iHS and XP-EHH statistics across the programs rehh (version 2.0), `selscan` and `hapbin`. Pairwise correlation for the iHS computed in the CEU and the JPT+CHB (in parenthesis) populations are given in the upper diagonal. Pairwise correlation for the XP-EHH computed across the CEU and JPT+CHB populations are given in the lower diagonal.

|  | REHH | **selscan** | **hapbin** |
| --- | --- | --- | --- |
| REHH | *na* | 0.991 (0.993) | 0.907 (0.945) |
| **selscan** | 0.985 | *na* | 0.907 (0.945) |
| **hapbin** | 0.986 | 0.994 | *na* |

## Conclusion

Although the R package rehh (Gautier & Vitalis, 2012) has been widely used since its first release, the increasing dimension of haplotype datasets typically available in most species led to serious limitations. This stimulated the development of alternative R-free solutions (Szpiech & Hernandez, 2014; Maclean *et al*, 2015). In this study, we introduced substantial changes in the rehh package to improve its computational efficiency by one to several orders of magnitude. This was achieved by modifying the processing of the input files and, most importantly, by improving and parallelizing the core algorithm that computes the distribution of haplotype counts. As a result, rehh 2.0 clearly outperforms the `selscan` (Szpiech & Hernandez, 2014) package and competes with `hapbin` (Maclean *et al*, 2015), the fastest program to date. A decisive advantage of rehh 2.0 over these programs is that it allows working within the multi-platform R environment. As such, it benefits from several graphical tools that facilitate visual interpretation of the results.

rehh 2.0 is available from the CRAN repository (http://cran.r-project.org/web/packages/rehh/index.html). A help file together with a detailed vignette manual (the current version is provided as a Supplementary File S2) are included in the package.

## Acknowledgment

We wish to thank all users of the previous version for their feedback that helped to improve the package. This work was supported in part by a grant of the German Science Foundation (DFG-SFB680) to AK.

## Supplementary Material

- File S1: compressed archive named `FileS1.tar.gz` containing example input haplotype data and SNP information files in the rehh, `selscan` and `hapbin` format. The archive also contains command lines that were used to run the different programs
- File S2: Detailed user manual (vignette) for the rehh 2.0.

